# AFM characterization of early *P. aeruginosa* aggregates highlights emergent mechanical properties

**DOI:** 10.1101/2025.06.30.662013

**Authors:** Caroline D. Miller, Meisam Asgari, Sophie E. Darch

## Abstract

*Pseudomonas aeruginosa* (*Pa*) is a leading cause of chronic lung infections in people with cystic fibrosis (pwCF), where its ability to form resilient, multicellular communities contributes to antibiotic tolerance and long-term persistence. While much of our understanding of *Pa* biofilms comes from surface-attached models, recent studies have emphasized the clinical relevance of suspended bacterial aggregates – dense, three-dimensional clusters that form early during infection and exhibit key biofilm-like properties. However, the physical characteristics of these aggregates remain poorly defined. Here, we apply atomic force microscopy (AFM) to visualize and quantify the structural and mechanical properties of *Pa* aggregates formed in synthetic cystic fibrosis sputum medium (SCFM2). Compared to planktonic cultures grown without mucin, aggregates formed in SCFM2 exhibited complex architecture and increased resistance to deformation, as measured by force spectroscopy. These differences emerged despite the absence of mature extracellular matrix components, suggesting that environmental cues and spatial organization alone may be sufficient to enhance aggregate mechanical resilience. Our results demonstrate that AFM provides a powerful, high-resolution approach for studying early-stage bacterial aggregates under physiologically relevant conditions. By resolving structural features and quantifying localized mechanical strength, this method offers new insight into how aggregate architecture contributes to persistence during chronic infection. These findings lay the groundwork for future studies targeting the physical robustness of bacterial communities as an early vulnerability in the pathogenesis of *Pa* both in CF and in other infection settings.

## Importance

Chronic infections in people with cystic fibrosis are notoriously difficult to treat, in part due to the ability of *Pseudomonas aeruginosa* (*Pa*) to form protective communities known as aggregates. These suspended, multicellular clusters are not well captured by traditional surface-attached biofilm models but are now recognized as an important feature of persistent infection. Understanding how these aggregates resist physical and antimicrobial disruption is essential for developing better therapies. This study uses atomic force microscopy (AFM) to examine *Pa* aggregates at nanometer resolution in a laboratory model that mimics cystic fibrosis (CF) lung secretions. AFM not only visualizes individual aggregates but also measures how strongly they resist being physically deformed. Our findings show that aggregates formed in this environment are structurally robust, compared to single cells. These results highlight the importance of early physical organization in bacterial persistence and suggest new directions for therapies aimed at disrupting bacterial communities before they become established.

## Observation

*Pseudomonas aeruginosa* (*Pa*) is a leading cause of chronic lung infections in people with cystic fibrosis (pwCF), where it establishes persistent, antibiotic-tolerant populations that are difficult to eradicate^1-3^. While surface-attached biofilms have long served as the dominant model for *Pa* persistence, recent studies have shifted focus to the formation and properties of smaller, suspended bacterial aggregates (∼10–1,000 cells)^4-8^. These aggregates represent a critical intermediate between planktonic cells and mature biofilms and exhibit key biofilm-like traits, including antibiotic tolerance and immune evasion. Despite their clinical relevance, the physical and mechanical properties of aggregates, particularly during early stages of formation, remain poorly understood, limiting our ability to define how they contribute to chronic infection.

Although bacterial aggregates have been linked to persistence and tolerance, methods for studying their structural and mechanical properties *in situ* remain limited. Traditional imaging approaches provide morphological insight but lack the resolution to probe the physical forces that govern cell-cell organization and material strength. To overcome this, we turned to Atomic Force Microscopy (AFM) – a powerful technique capable of simultaneously capturing high-resolution images and quantifying nanoscale mechanical properties of bacterial communities^9-11^.

AFM offers unique insight into the physical properties of bacterial surfaces (including lipopolysaccharides (LPS) and extracellular polymeric substances (EPS))^12-15^. It has been instrumental in quantifying adhesion forces between bacteria and surfaces, as well as intercellular interactions within communities - key parameters for understanding biofilm formation and stability^9, 16-20^. Previous AFM studies have shown that *Pa* strains with distinct LPS profiles exhibit different physical properties that also influence their capacity to form biofilms^5, 12, 14^. The technique has also revealed the contributions of biofilm-associated structures such as type IV pili, Pel, Psl, and extracellular DNA (eDNA) to biofilm architecture and cohesion^12, 14, 16, 17^. However, these analyses have largely focused on mature, surface-attached biofilms. By shifting attention to suspended aggregates, we gain higher resolution into early-stage organization. Studying smaller aggregates allows for single-cluster and even single-cell level analysis – offering a more detailed view of the rapid aggregation processes that occur within hours of infection. This approach sheds light on the immediate physiological and mechanical changes that accompany the transition from free-living cells to organized multicellular communities.

Here, we applied AFM to visualize and measure the physical properties of *Pa* aggregates formed in synthetic cystic fibrosis sputum medium (SCFM2), a clinically relevant *in vitro* model that mimics the biochemical and rheological properties of the CF lung environment. By comparing aggregates formed in SCFM2 to planktonic cells grown in the absence of mucin, we aimed to characterize structural and mechanical features that might contribute to aggregate persistence. AFM is uniquely suited to probe bacterial surfaces and interactions at nanometer resolution. Beyond high-resolution imaging, AFM enables force mapping and nanoindentation, allowing us to quantify the resistance of individual bacterial aggregates to deformation. These localized mechanical measurements offer insight into the internal organization and robustness of aggregates without disrupting their native state.

In this study, we bridge the gap between planktonic and biofilm states by using atomic force microscopy to examine the structure and mechanical strength of *Pa* aggregates formed in a synthetic cystic fibrosis sputum medium. This approach provides a proof of concept for high-resolution, *in situ* analysis of bacterial communities in CF-like conditions. By revealing physical features that distinguish aggregates from planktonic cells, our findings offer mechanistic insight into early-stage resilience and may inform new strategies to disrupt bacterial persistence before mature biofilms form.

### Mucin promotes aggregate formation and complex architecture in CF-like conditions

We cultured wild-type *P. aeruginosa* (*Pa)* (PAO1::pMRP9-1 (GFP)) in synthetic cystic fibrosis sputum medium (SCFM2) with and without mucin under static conditions to assess early-stage aggregate development.

Cultures supplemented with mucin consistently produced dense, spatially distinct bacterial aggregates, while cultures lacking mucin yielded predominantly dispersed, planktonic cells. These observations are consistent with previous work demonstrating that mucin facilitates bacterial clustering in our modelled CF lung environment, resulting in the formation of aggregates of similar sizes to those observed *in vivo*^4^. After 4 hours of growth, cultures were gently transferred onto poly-L-lysine-coated glass slides to enable surface attachment for imaging and force spectroscopy. This workflow, illustrated in our schematic (Figure 1), preserves the native structure of aggregates while allowing for high-resolution AFM analysis. AFM imaging of aggregates formed in SCFM2 revealed complex topographical features. The aggregates displayed a multilayered structure with tightly packed cells and variable surface elevations. In contrast, planktonic cells in mucin-free conditions appeared as isolated, smooth, rounded morphologies with little interaction between neighboring cells. These findings indicate that mucin not only promotes aggregate formation but also drives structural reorganization that resembles the architecture of early-stage biofilms, even in the absence of surface attachment.

**Figure 1.**
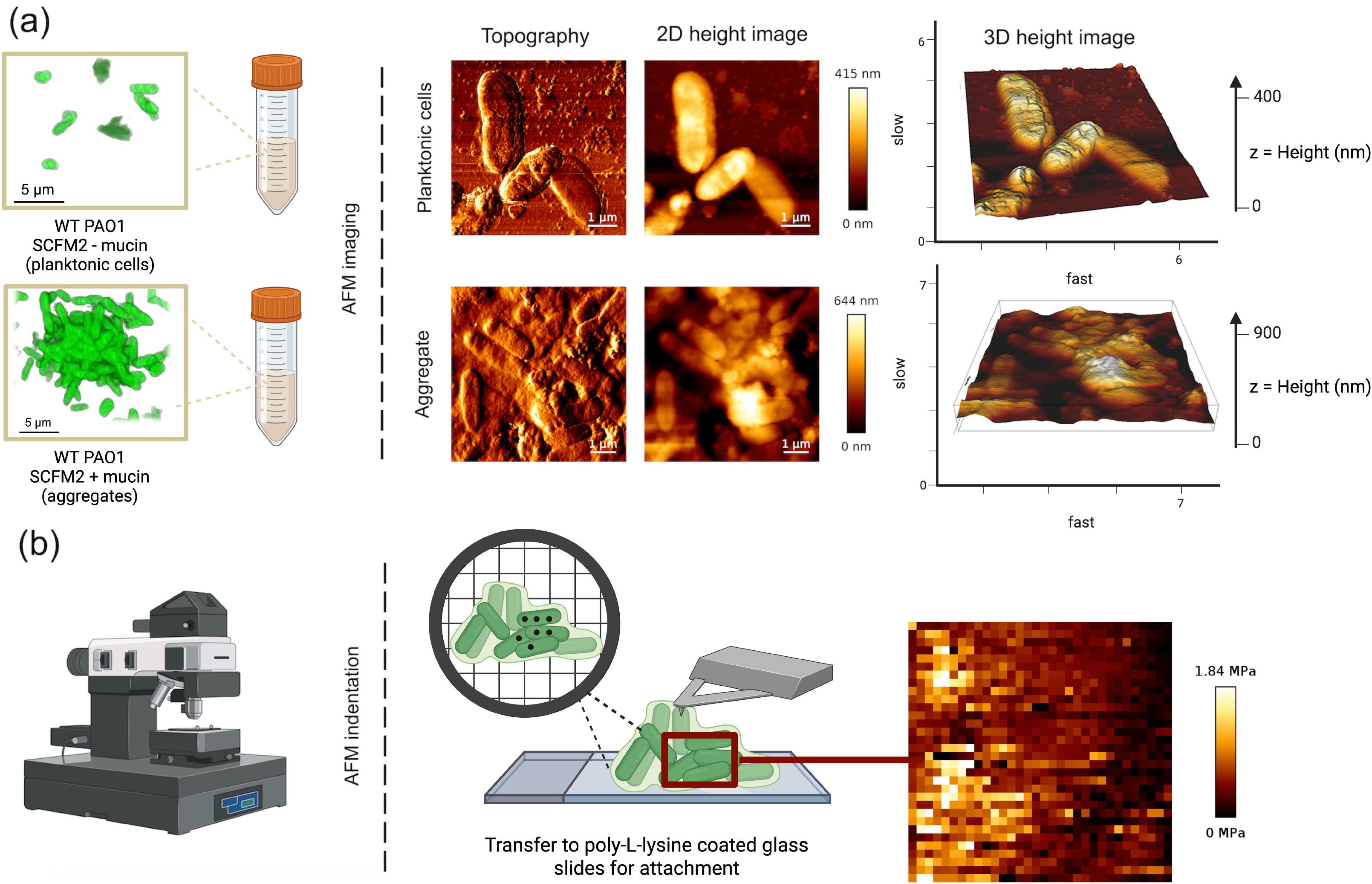
Assessing *P. aeruginosa* aggregate formation using Atomic Force Microscopy (AFM). **(a)** Schematic of methods used for *P. aeruginosa* PAO1 culture in SCFM2 (with or without mucin) to model aggregate formation within the CF lung environment or planktonic growth, respectively. PAO1 is cultured in synthetic CF sputum to produce planktonic cells or aggregates, as represented by 3D renderings of confocal laser scanning microscopy images (scale bar 5 μm). Samples are transferred to poly-L-lysine coated glass slides for attachment and AFM is used to extract topographical 2D and 3D imaging data. **(b)** Example shows measurement of physical forces using localized force spectroscopy to quantify the elastic modulus (measure of stiffness, MPa) and resistance to deformation at multiple positions across a single cell.

### Aggregates exhibit increased mechanical strength relative to planktonic cells

To quantify the physical differences between aggregates and planktonic *P. aeruginosa*, we used localized AFM force spectroscopy to measure the elastic modulus of the samples. Aggregates formed in SCFM2 with mucin exhibited significantly higher mechanical stiffness than planktonic cells grown in mucin-free media. Specifically, the average elastic modulus of aggregate regions was approximately mean: 218.7 ± 118.7 kPa, n *=* 2,843 compared to mean: 50.8 ± 35.8 kPa, n = 3,915 for individual planktonic cells (n = number of individual indentation measurements per condition). A two-tailed unpaired t-test confirmed this difference was highly significant (*t* = 73.07, *p* < 0.0001). These elastic modulus values are intuitively consistent with the representative force–distance curves shown in Fig. 2a, which demonstrate penetration forces between the two cell types differ while both are on the order of 0.3–0.4 nN. As shown in Figure 2, adhesion was negligible due to the use of a large spherical tip, so the Hertzian model was applied, consistent with prior work on low-adhesion biological systems^21-23^.A s expected for AFM indentation of live bacteria, individual force curves showed substantial variability, contributing to standard deviations approaching ∼50% of the mean. While certain indentation curves reflect standard deformation without compromising the cell membrane, others exhibit signs of membrane perforation due to localized force fluctuations, as illustrated in Figure 2. Nonetheless, aggregates consistently resisted indentation more strongly than planktonic cells, supporting the conclusion that early-stage multicellular structure alone confers increased mechanical integrity.

**Figure 2:**
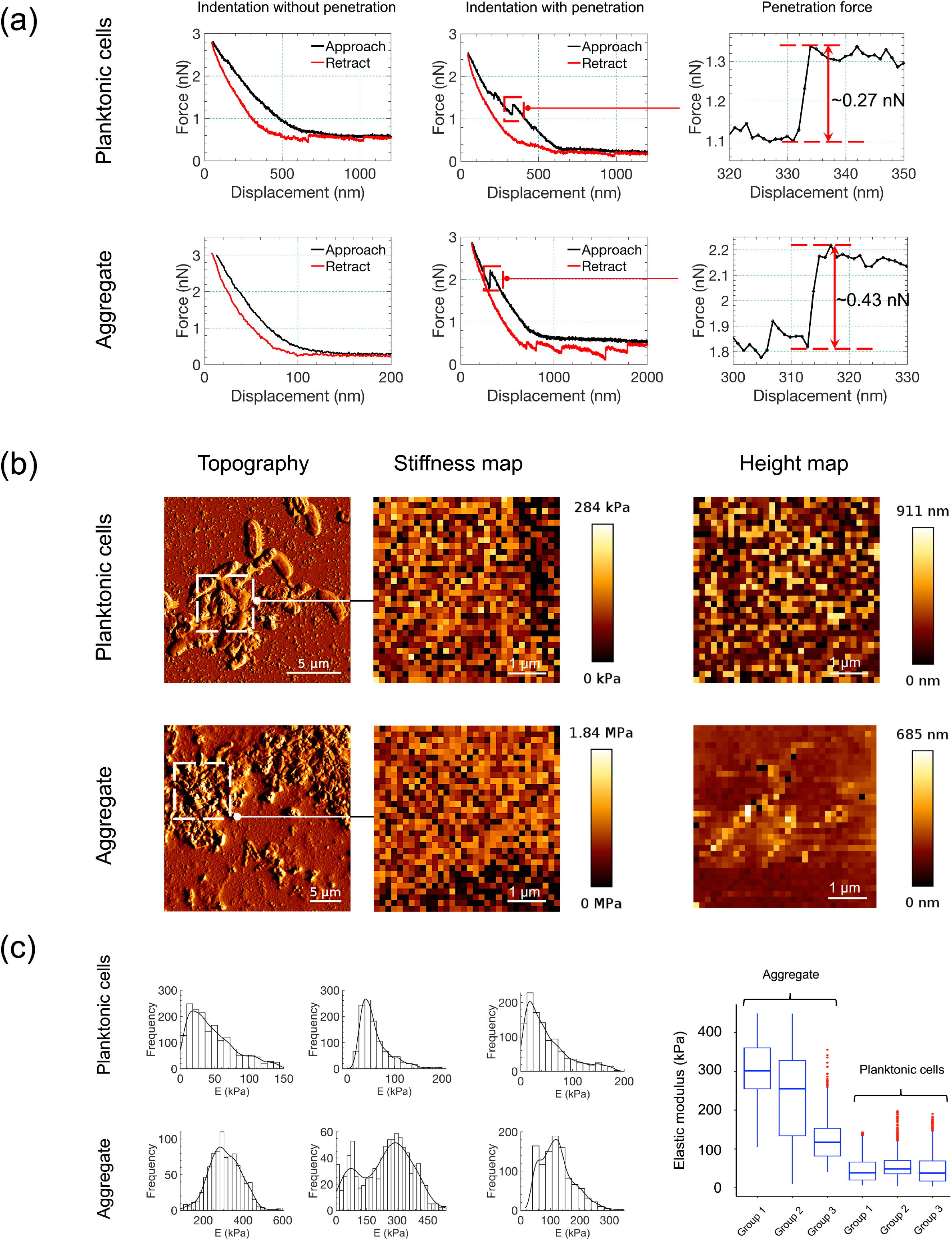
Atomic force microscopy (AFM) reveals increased stiffness and structural complexity in early *P. aeruginosa aggregates*. **(a)**Representative force-displacement curves comparing planktonic cells (SCFM2 without mucin) and aggregates (SCFM2 with mucin), captured by AFM using spherical-tipped CONT-Au cantilevers. Curves illustrate indentation with and without penetration, and calculated penetration forces for select interactions. Data represents 3 biological replicates of each condition (+/-mucin), containing ∼2-4,000 measurement points. **(b)**Topography, elastic modulus, and height maps derived from AFM imaging and force spectroscopy. Aggregates formed in SCFM2 exhibit greater mechanical heterogeneity, increased stiffness (up to 1.84 MPa), and multilayered structure compared to dispersed planktonic cells. Data represents 3 biological replicates of each condition (+/-mucin), containing ∼2-4,000 measurement points. Scale bars are 5 μm, 1 μm and 1 μm for topography, elastic modulus and height maps respectively. **(c)**Histograms of localized elastic modulus values extracted from six independent regions per condition. Aggregates show a broader and right-shifted modulus distribution relative to planktonic cells, indicating significantly increased stiffness (p < 0.0001, unpaired t-test). Box plots represent elastic modulus data for 3biological replicates of each sample type (planktonic cells or aggregates). All modulus values were calculated using the Hertz contact model.

This enhanced mechanical strength emerged despite the absence of mature exopolysaccharide scaffolding, indicating that cellular reorganization and compaction within a mucus-rich environment is sufficient to confer resilience. Transcriptomic analyses confirm that *pel, psl*, and *alg* genes are not expressed at this early stage^24^, and AFM imaging similarly shows no evidence of exopolysaccharide structures. Instead, the tightly packed, layered architecture observed in aggregate AFM images likely contributes to this mechanical robustness.

Such reinforcement may represent an early physical adaptation that protects bacteria from shear stress, host immune mechanisms, and antimicrobial exposure, even before significant deposition of matrix components traditionally associated with biofilm formation. While these measurements were performed on poly-L-lysine– coated glass, which is stiffer than lung tissue, this approach provided the stability required for reproducible nanoscale interrogation of single aggregates and represents a critical first step toward linking aggregate architecture with biomechanical resilience.

Together, these data support a model in which *Pa* aggregates rapidly acquire distinct biomechanical properties that differentiate them from planktonic cells. These structural adaptations may underlie the observed tolerance of aggregates *in vivo* and reinforce the importance of targeting aggregate formation at early stages of infection. AFM enables *in situ* interrogation of these properties with nanoscale resolution, offering a powerful platform for dissecting how host factors, bacterial genetics, and environmental cues shape the physical resilience of bacterial communities.

## Conclusion

These findings underscore the potential clinical relevance of targeting early aggregate formation before mature biofilms develop. By identifying that *Pa* aggregates rapidly acquire mechanical properties that may contribute to persistence, our study highlights a window of vulnerability that could be exploited by novel therapeutic strategies. Future work will focus on how specific genetic or environmental factors modulate aggregate mechanics, and whether these properties correlate with antibiotic tolerance or immune evasion. Ultimately, integrating biophysical profiling into infection models could inform more effective, stage-specific interventions for managing chronic *Pa* infections in pwCF and other at-risk populations.

## Acknowledgements

S.E.D is supported by start-up funds provided by the Department of Molecular Medicine, The University of South Florida, as well as research grants from the Cystic Fibrosis Foundation (CFF) (DARCH19G0, DARCH22P0). C.D.M is supported by a CFF student award (MILLER24H0).

## Author Contributions

S.E.D. and C.D.M. conceived the project, designed experiments, performed data analysis, prepared figures, and wrote the manuscript. C.D.M. performed the bacterial experiments. M.A. performed atomic force microscopy (AFM) experiments, conducted AFM data analysis, and prepared the AFM figures.

---

**Figure S1:**
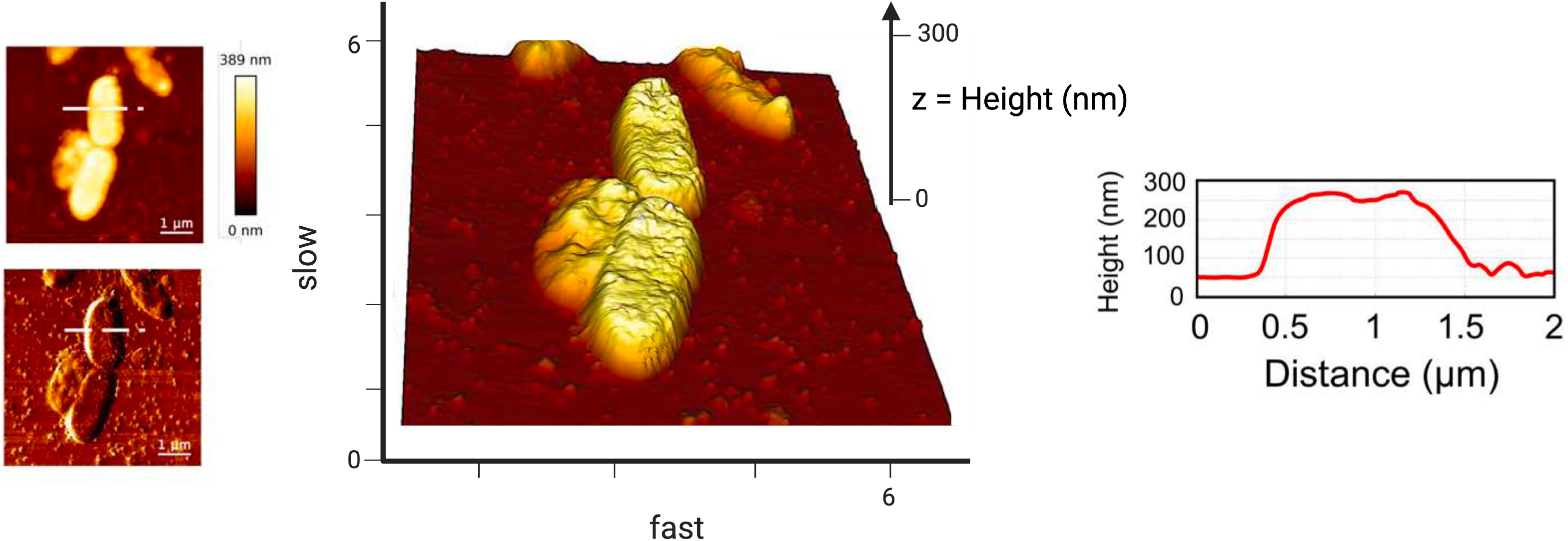
AFM cross-sectional height profile of *Pseudomonas aeruginosa* within an aggregate. Representative AFM height images and corresponding cross-sectional analysis demonstrate the vertical dimension of individual *P. aeruginosa* cells within an early-stage aggregate (4 hours growth). The 3D topography (center) and line profile (right) indicate an average cell height of ∼300 nm, consistent with previously reported dimensions of hydrated *P. aeruginosa* cells in aggregates. Additional AFM height maps (left) show top-view scans with cross-section lines used for profiling. Together, these images confirm that aggregates represent physiological *in-situ* dimensions during AFM imaging and analysis.

## Supplementary Information (SI)

### Materials and methods

#### Bacterial strains and growth conditions

*Pseudomonas aeruginosa* PAO1 wild type carrying the plasmid pMRP9-1, expressing GFP, was cultured in standard lab media (LB) from frozen stock overnight at 37°C with shaking (200 rpm)^1, 2^. Cells were back diluted 1:20 in fresh LB and grown to log-phase (∼3 hours), then washed with PBS (pH7.0) before inoculation^2^. For aggregates, SCFM2 containing mucin (SCFM2 + mucin) (Porcine mucin, 250 mg) was prepared as previously described^2, 3^ (obtained from SynthBiome). For planktonic cultures, SCFM2 without mucin (SCFM2 - mucin) was prepared identically but with the exclusion of mucin^2^. Washed *Pa* cells were inoculated into both media at approximately 10^5^ cells per mL (0.05 OD^600^) and incubated without shaking at 37°C for 4 hours to allow for aggregate formation^2, 4^.

#### Atomic Force Microscopy (AFM) analysis of *P. aeruginosa* aggregates and planktonic cells

#### Sample preparation

For both growth conditions (SCFM2 -/+ mucin), 4-hour cultures were diluted 1:20 in PBS to create a ‘low density culture’. A hydrophobic pen was used to create a liquid repellent area (circular) on poly-L-lysine coated microscope slides (Fisher Scientific) to which 20 μL of sample was pipetted into the center.

Throughout sessions, 100-200 μL of molecular grade water was supplemented as needed for hydration and to improve resolution of individual cells and/or aggregates.

#### Atomic Force Microscopy

A JPK Nanowizard PURE atomic force microscope (Bruker Nano, Berlin, Germany) mounted on an inverted epifluorescence Zeiss Axiovert 200 M microscope (Carl Zeiss Microscopy, Göttingen, Germany) was utilized for imaging and mechanical characterization of *P. aeruginosa* samples at the micro-to nanoscale. The samples were imaged and mechanically tested following the developed protocols^5^.

Multiple images were captured at each location on each sample using contact mode imaging. The maximum lateral scan focused on regions measuring 50 μm × 50 μm, with a scanning rate set to 1 Hz. Reduced scan areas were then selected to obtain detailed structural information of the samples. MSNL-10 silicon nitride cantilevers (Bruker, Mannheim, Germany) with a spring constant of 0.01-0.1 N/m and a nominal tip radius of 2 nm were used for imaging. To ensure that the same region of interest was analyzed following probe exchange, we used the motorized XY stage of the JPK Nanowizard system. The stage allows precise, computer-controlled repositioning to previously defined coordinates, which are stored by the software during the initial high-resolution scan. After changing to the blunt probe for force spectroscopy, the stage was guided back to the exact same coordinates. To confirm correct alignment, topographical landmarks within the selected region were cross-checked before indentation measurements were performed. This approach ensured reproducibility of measurements from the same imaged area without loss of positional accuracy. For indentation measurements, Biosphere Au Reflex (CONT-Au) cantilevers (Nanotools USA LLC, Henderson, NV) with a nominal spring constant of 0.2 N/m, a length of 450 μm, and a nominal resonance frequency of 13 kHz in air, equipped with integrated spherical tips of radii 100 nm and 2 μm (±10%), were employed ^6, 7^. All measurements were conducted in a wet environment with the sample immersed in PBS. The indentation rate was set to 2 μm/s. Before each test, the deflection sensitivity of the cantilever was calibrated by engaging it on a clean glass slide in a wet state ^6, 7^. The exact spring constant of the cantilever was calibrated using thermal noise fluctuations in air ^5^. This was done by fitting the first free resonance peak of the cantilever to that of a simple harmonic oscillator using the JPK software ^5^. The software employs various models, such as Hertz and Sneddon contact mechanics, to extract the elastic modulus from force-displacement curves obtained during indentation tests ^8^. The elastic modulus was obtained from the Hertzian contact model ^8^. In this model, the contact radius a is related to the indenting force F through

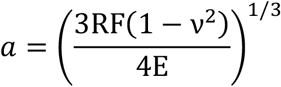

with R being the radius of the spherical tip, and ν and E being the Poisson’s ratio and elastic modulus of the sample, respectively ^8^. The indentation depth δ is expressed in terms of the contact radius as

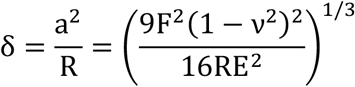

The JPK data processing software was used to analyze the indentation data.

